# Quantitative trait loci (QTL) underlying phenotypic variation in bioethanol-related processes in *Neurospora crassa*

**DOI:** 10.1101/736470

**Authors:** Joshua C. Waters, Deval Jhaveri, Justin C. Biffinger, Kwangwon Lee

## Abstract

Bioethanol production from lignocellulosic biomass has received increasing attention over the past decade. Many attempts have been made to reduce the cost of bioethanol production by combining the separate steps of the process into a single-step process known as consolidated bioprocessing. This requires identification of organisms that can efficiently decompose lignocellulose to simple sugars and ferment the pentose and hexose sugars liberated to ethanol. There have been many attempts in engineering laboratory strains by adding new genes or modifying genes to expand the capacity of an industrial microorganism. There has been less attention in improving bioethanol-related processes utilizing natural variation existing in the natural ecotypes. In this study, we sought to identify genomic loci contributing to variation in saccharification of cellulose and fermentation of glucose in the fermenting cellulolytic fungus *Neurospora crassa* through quantitative trait loci (QTL) analysis. We identified one major QTL contributing to fermentation of glucose and multiple putative QTL’s underlying saccharification. Understanding the natural variation of the major QTL gene would provide new insights in developing industrial microbes for bioethanol production.

## Introduction

Fermentation of cellulosic biomass by microorganisms is a complex process requiring coordinated regulation of multiple pathways, including production and secretion of at least three distinct classes of catabolic enzymes (cellulase, hemicellulase, and lignin peroxidase enzymes), transport of substrates into the cell, followed by catabolism and fermentation of the resultant monomeric C5 and C6 sugars. Identification of organisms efficient in all requisite processes has been a major limitation toward realizing the vision of the consolidated bioprocessing (CBP) of biomass to ethanol for fuels (1–4). While many researchers attempt to engineer existing strains by introducing the genes for pathways lacking in the given organisms, success has been limited due to the increased physiological burdens imposed by genetic manipulation. There is also a growing concern that genetic manipulations that are effective in laboratory strains may not be translatable to more robust strains used in industry (3, 5). There are, however, organisms that already possess the genetic elements needed to perform all of the requisite processes, such as the model filamentous fungus *Neurospora crassa* [9].

These types of native microorganisms present excellent potential sources of useful industrial enzymes and products. Breeding has proven to be a reliable method for manipulating genomes to select for desired traits, as evidenced by the successes of plant breeding (6). Selective mating has long been used in plant breeding to improve traits ranging from fruit size and yield to appearance or flavor (7, 8). Recently, selective breeding has also been used in fungi in attempts to select for increased saccharification and fermentation potential (9).

The advent of computational tools such as genome-wide association studies (GWAS) and quantitative trait loci (QTL) analysis have had drastic impacts on how plant geneticists design new lineages with improved traits and qualities (10). GWAS and QTL analysis are computational tools for investigating the genetic elements underlying quantitative traits. Unlike qualitative traits such as color, which are inherited discretely and exhibit discontinuous variation, quantitative traits exhibit continuous variation and cannot be discretely grouped (10). GWAS and QTL use polymorphic genetic markers to predict the extent to which a given genetic locus contributes to observed variation in quantitative traits, such as plant height, and researchers have been able to use genetic methods to identify combinations of desired alleles in individuals that are then used for breeding, known as marker assisted selection (MAS).

QTL analysis and MAS have revolutionized plant genetics and should be equally useful for improvement of quantitative traits in fungi, such as saccharification and fermentation of cellulose. While a wealth of information is available on the effects of gene deletions or overexpression of cellulolytic enzymes or fermentation pathways, there have been few studies to characterize the variation resulting from allelic variation in populations, most of which focus solely on fermentation (11–18, 18–21). While gene deletions and overexpression studies can implicate genetic elements in these processes, these manipulations increase cellular burden due to unknown pleiotropic effects, and multiple manipulations can result in compounding deficits (3, 5, 22, 23).

The inherent complexity of lignocellulose metabolism allows for abundant sources of variation that will affect CBP performance. Understanding how allelic variation attenuates performance will lead to a more robust understanding of cellulose metabolism and identify superior combinations of alleles for trait enhancement for CBP. To this end, we chose to perform QTL analysis of bioethanol-related processes, specifically saccharification and fermentation, on a laboratory generated population of the model filamentous fungus *Neurospora crassa*. 111 offspring and two parental strains were genotyped by sequencing, evaluated for their ability to decompose cellulose and ferment glucose, and subjected to QTL analysis. A major QTL was identified for fermentation and multiple putative QTL’s were identified for saccharification of cellulose.

## Materials and methods

### Strains

Parental strains FGSC2223 (Mat a, collected in Iowa, USA) and FGSC4825 (Mat A, collected in Ivory Coast, Africa) from the Fungal Genetics Stock Center (FGSC) (24), and their cross progeny (designated as the N6 population) were grown from long-term stock prior to each experiment on Vogel’s minimal media slants (1X Vogel’s Salts, 2% Sucrose, 1.5% Agar, pH 5.8). Spore suspensions in High Glucose Liquid Media (HGLM) (1X Vogel’s salts, 2% glucose, 0.5% L-Arginine, pH 5.8) were used to generate mycelial mats in petri plates, which were used to generate replicate mycelial pads for each experiment using a bore punch.

### Enzyme activity assay

A modified FPA assay was performed using 96-well plates as described by Camassola and Dillon, in which secreted protein extracts were taken from wells in 6-well plates containing 1% CMC broth (1X Vogel’s salts, 1% CMC) 4 days after inoculation with 10 gauge mycelial pads (25). Before inoculating the mycelia pads, they were rinsed in sterile water three times to remove media. Culture broth containing secreted enzyme was filtered and centrifuged at 13.2k rpm to remove any fungal cells or debris. 50μL of supernatant was added to 100μL of 50mM sodium acetate buffer pH 5.6 in a 96-well deep-well plate, which was then equilibrated to 50°C for 5 min in a hot-water bath. A 5mg strip of Whatman Grade No. 1 filter paper was submerged in the solution, and the plate was incubated at 50°C for 60 min. After 60 min, 300μL of 3,5-dinitrosalicylic acid (DNS) Reagent was added to stop the enzymatic reaction and visualize glucose equivalents. The plate was incubated at 100°C for 10 min to develop color, then transferred to an ice bath to stop color development. 100μL of Enzyme/DNS mixture was transferred to a clear-bottom assay plate, diluted with 200μL of deionized H_2_O, and absorbance was measured at 545 nm. The concentration of reducing equivalents released was determined with standard curves generated with glucose standards of known concentration. All samples were performed in biological triplicates. The activity could not be quantified in conventional Filter Paper Units because of the low concentration of secreted enzymes. Therefore, these results were presented in the form of the concentration of reducing equivalents released in each aliquot.

### Fermentations and ethanol quantification

To characterize fermentation of glucose among the lab generated N6 first filial (F1) generation and the parental strains (FGSC 2223 and FGSC4825), fermentation was carried out in a 96-well format in deep-well plates. Biological replicates for each strain were collected from mycelial mats grown from spore suspension in high glucose liquid media (HGLM) with a 6 gauge punch and inoculated into 750μl of HGLM (2% glucose), sealed with aluminum Thermowell^™^ seals, and allowed to ferment for 7 days in 12:12 Light/Dark conditions at 25°C. All samples were performed in biological quadruplicate. After fermentation, 600 μL of media was recovered and cell debris was removed by sequential centrifugation at 13.2k rpm for 5 min. The recovered supernatant was aliquoted into HPLC vials for ethanol analysis by HPLC. HPLC quantitation was performed using a Varian ProStar HPLC with a Varian ProStar Autosampler and a Varian 356-LC Refractive Index Detector. An isocratic elution was used with an Agilent Hi-Plex H^+^ (300 mm × 7.7 mm ion-exchange column) with 5mM H_2_SO_4_ at a flow rate of 0.7 ml/min at 60°C. The concentration of ethanol present was determined from a standard curve based on ethanol standards with a known concentration.

### Qualitative Trait Loci (QTL) Analysis

To investigate the underlying sources of variation in saccharification and fermentation, a qualitative trait loci (QTL) analysis was performed using R to identify candidate genes contributing to the observed variation. N6 population has been used for QTL analysis using simple sequence repeat markers. For the current study, we generated a single nucleotide polymorphism (SNP) library using the Genotyping By Sequencing (GBS) approach (26). GBS data for the current study are available at (SRA accession: PRJNA594422.). Briefly, 50,000 SNP markers were filtered to 146 evenly distributed informative markers using Python and later Excel. First, all markers that were not polymorphic between parents were removed, followed by any markers in which one of the two parents was missing data. The resulting 4900 markers were formatted for Excel for further filtering. Chi-square tests were performed for each marker and those markers with unequal segregation among progeny (>20% disparity) were removed, followed by markers with >10% missing data among progeny. The filtered genotype data was combined with phenotype data from FPA and glucose fermentation assays and formatted for R-QTL. The formatted data was imported into R-QTL as and the create map function was used to generate a linkage map from a physical map based on recombination frequencies. Markers were then hand curated to generate a linkage map for all 7 chromosomes (Linkage groups) of *Neurospora crassa* with approximately 22 evenly spaced markers per chromosome.

After generating a linkage map for the N6 population, QTL scans were performed for each phenotype. Composite interval mapping (CIM) was used to scan for single QTL’s using the maximum likelihood (em), Haley-Knott regression (hk), and multiple imputation (imp) methods in the R-QTL package, allowing for up to 4 covariate markers within a 20-marker window. Likelihood odds ratio (LOD) thresholds were generated for 95% and 90% confidence intervals for each method based on 1000 permutations of the data. Putative QTL were designated as major QTL if they explained greater than 10% of the observed variation (27). Peak marker positions for each chromosome (based on LOD score) were used to identify linkage blocks between flanking markers to identify chromosomal regions containing the suspected QTL, according to their physical positions. These chromosomal regions were searched in the Fungi Database (FungiDB.org) to identify all genes within the region to find candidate genes contributing to the observed variation.

## Results

To test if there exist a natural variation of the bioethanol-related processes in one mapping population N6, we performed a QTL analyses in N6. Significant variation was observed among the N6 population for both saccharification of cellulose and fermentation of glucose (Fig 1). Both traits demonstrated transgressive segregation, with 21 or 58 of 111 offspring outperforming the more proficient parent and 66 or 33 of 111 underperforming the lesser parent in the FPA assay or fermentation of glucose, respectively (t-test, p = 0.00232, 0.0340).

**Figure 1.**
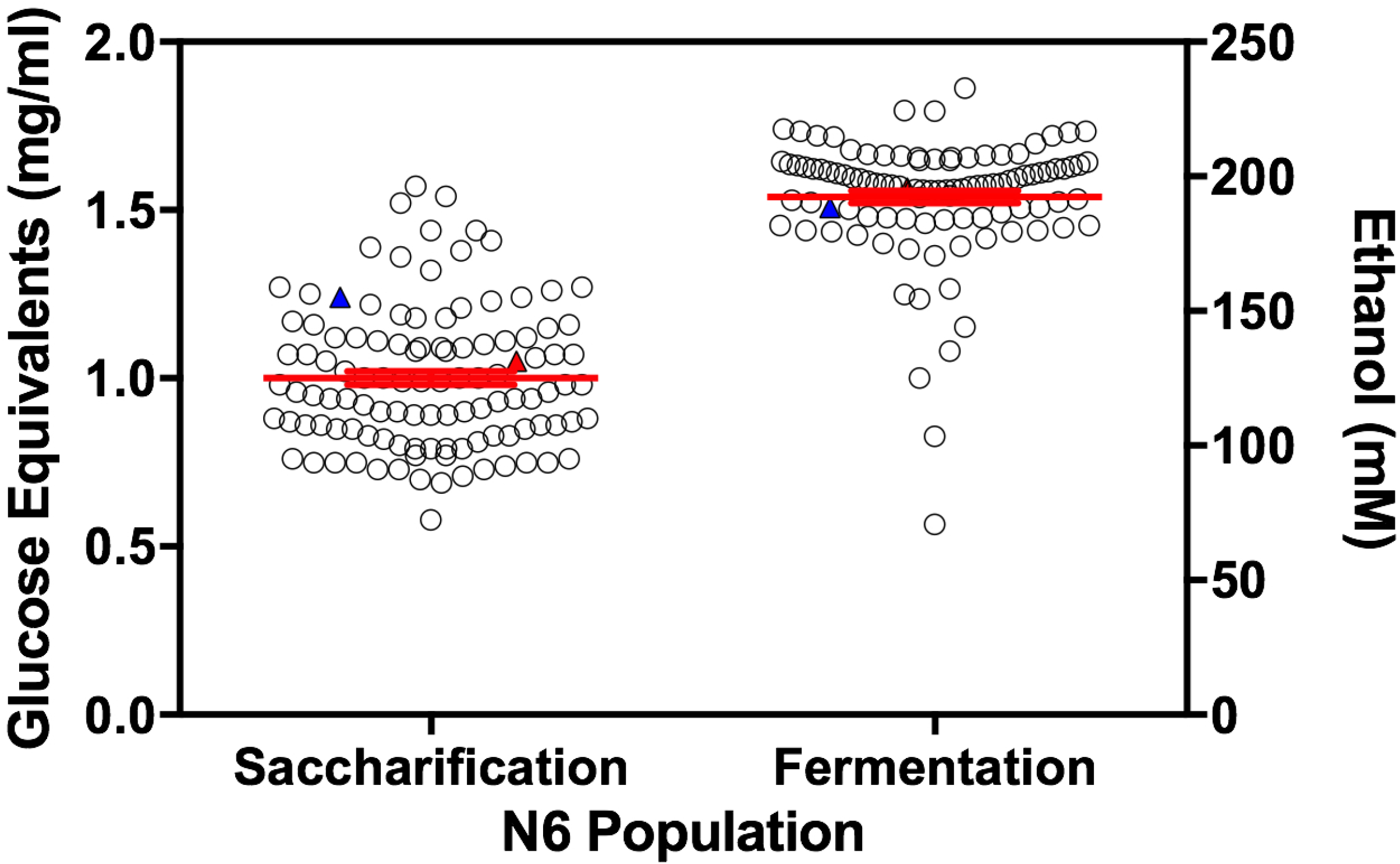
Natural variation in cellulolytic activity and fermentation among the N6 population. Significant variation was observed for saccharification of cellulose (left axis) and fermentation of glucose (right axis) among the N6 population (ANOVA p=3.99×10^-33^ and p=8.75×10^-17^ respectively). Each data point represents the mean of 3 biological replicates. Red bars represent mean with SEM. Red filled triangle represents FGSC4825. Blue filled triangle represents FGSC2223.

Due to the low level of recombination from only 1 meiotic event in a single cross, generating a map with evenly spaced markers required filtering a large number of polymorphic markers, reducing the power of QTL analysis. A linkage map based on recombination frequencies was generated from a physical map of SNP markers with approximately 22 evenly spaced markers on each of *N. crassa’s* 7 chromosomes (Fig 2).

**Figure 2.**
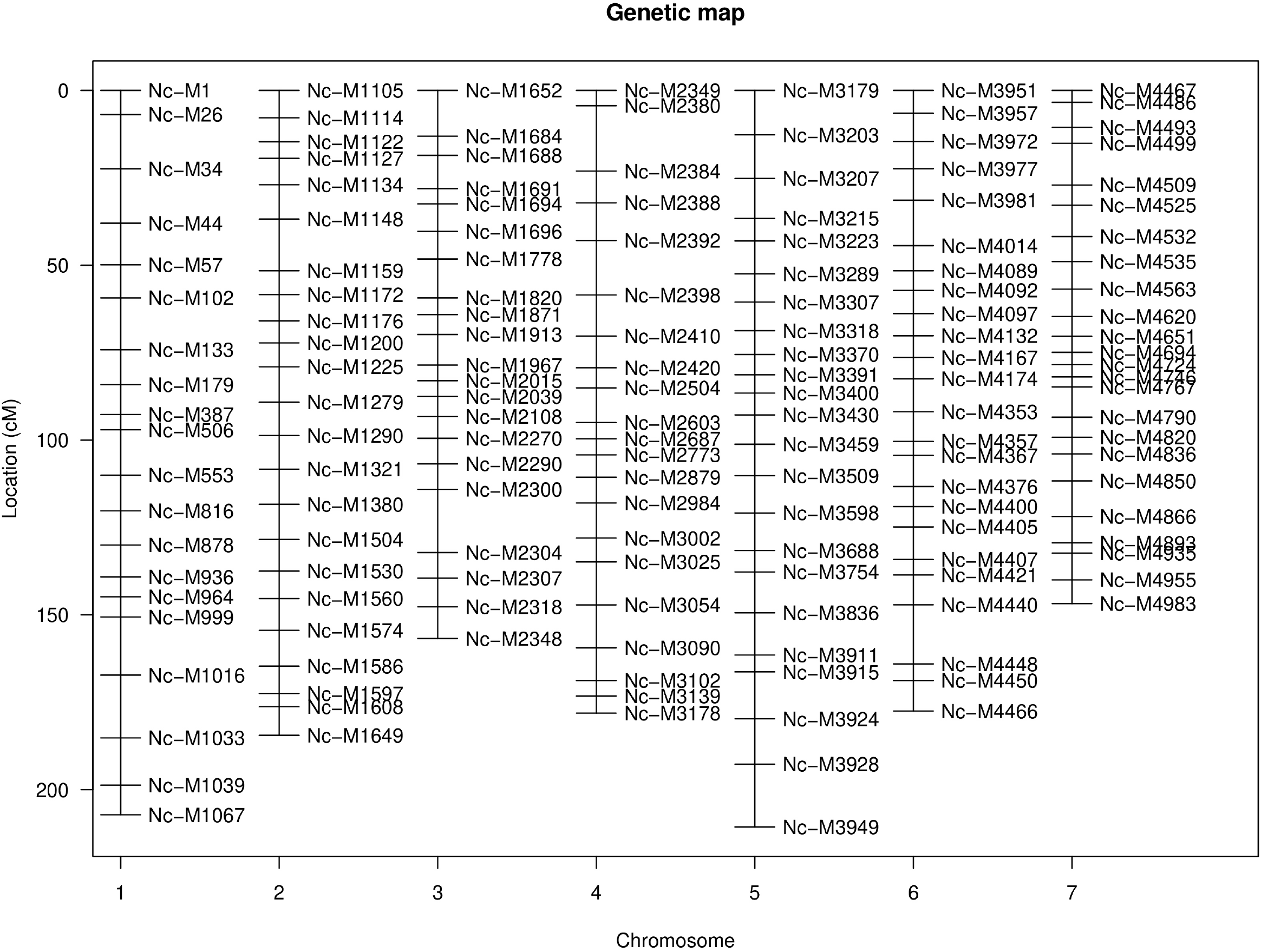
Linkage map of SNP markers for the N6 population. A linkage map created in R-QTL from physical marker positions, with approximately 22 evenly distributed markers per chromosome. Locations and distance are measured in centimorgans (cM).

A major QTL region contributing to variation in fermentation was identified on Linkage Group (LG) I (Nc-M878 = CM002236: 7449359: 361 Genes - 160 Annotated, 142 Hypothetical, 59 Unspecified) within a 90% confidence interval (CI) (p-value=0.075) (Fig 3). Additionally, minor QTL’s were observed for fermentation on LG IV (Nc-M2879 = CM002239: 4344007: 344 Genes - 158 Annotated, 134 Hypothetical, 52 Unspecified), LG VI (Nc-M4092 = CM002241: 1218567: 62 Genes - 28 Annotated, 32 Hypothetical, 2 Unspecified), and LG VII (Nc-M4866 = CM002236: 1218567: 77 Genes - 30 Annotated, 37 Hypothetical, 10 Unspecified), however, none of these were above the LOD threshold for 90% CI. Marker Nc-M878 on LG I lies within DUF1212 domain membrane protein (NCU00717). Markers Nc-M2879 on LG IV and Nc-M4092 on LG VI lie within hypothetical proteins (NCU07332 and NCU05646 respectively), while marker Nc-M4866 on LG VII lies within an intergenic region.

**Figure 3.**
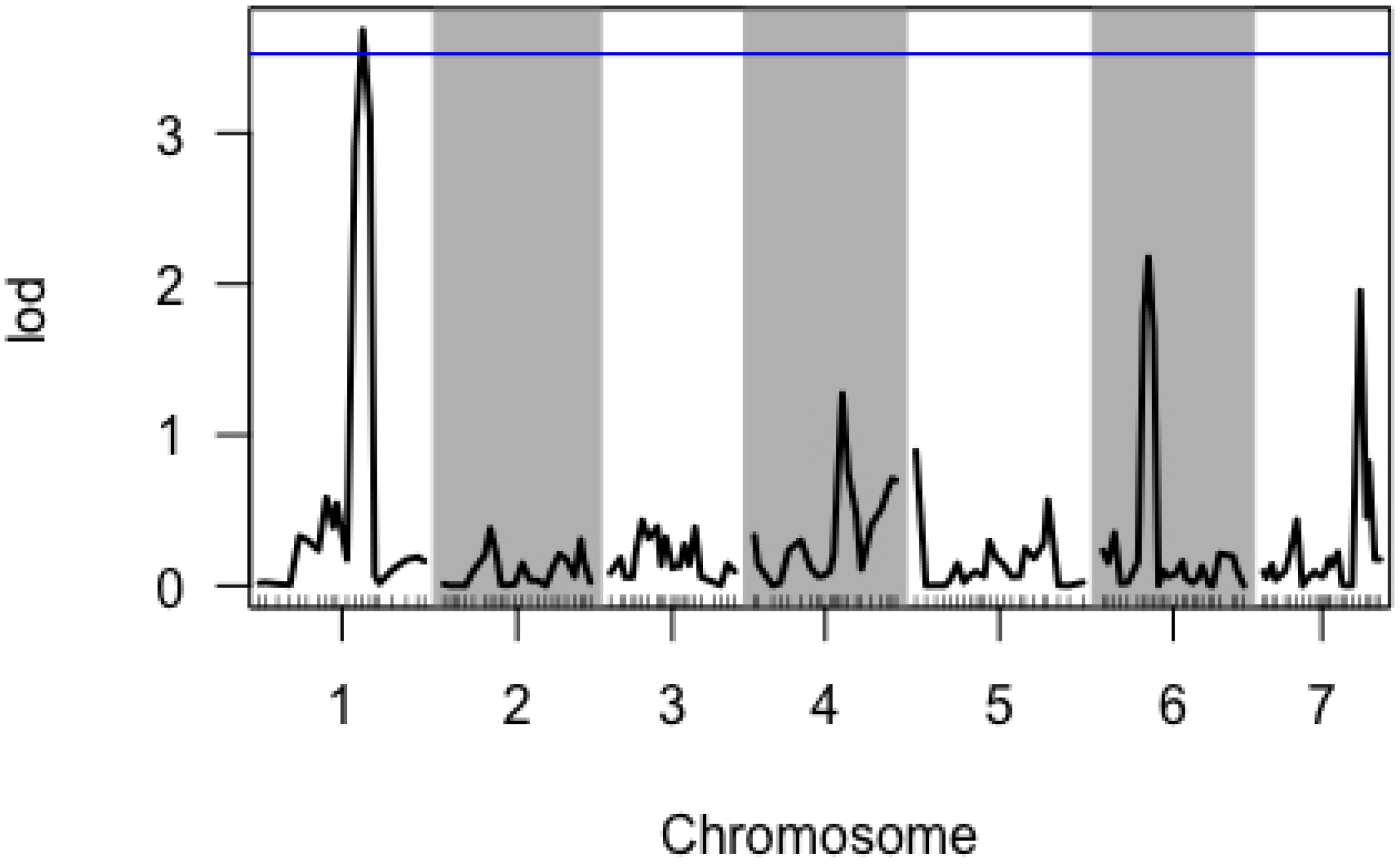
QTL analysis of Fermentation of Glucose to Ethanol by the N6 population. Composite Interval Mapping single-QTL scan for fermentation of glucose using linear regression (EM Algorithm). The y-axis is a measure of the logarithm of the odds ratio (LOD). Tick marks along the x-axis delineate relative marker positions, while alternating grey bands denote different chromosomes Blue line represents 90% confidence interval.

Using the fitqtl function, it was observed that the 4 QTL markers account for ~26% of the observed variation in fermentation of ethanol. The QTL on LG I accounts for 13% of the observed variation, while QTL on LG IV, VI and VII account for 4, 7, and 6% of the variation, respectively. Interestingly, marker segregation analysis revealed that the parental genotypes contributing to greater ethanol production varied between markers, with parental genotype A accounting for higher production at marker Nc-M878 on LG I and genotype B accounting for higher production at marker Nc-M4092 on LG VI (Fig 4A and B).

**Figure 4.**
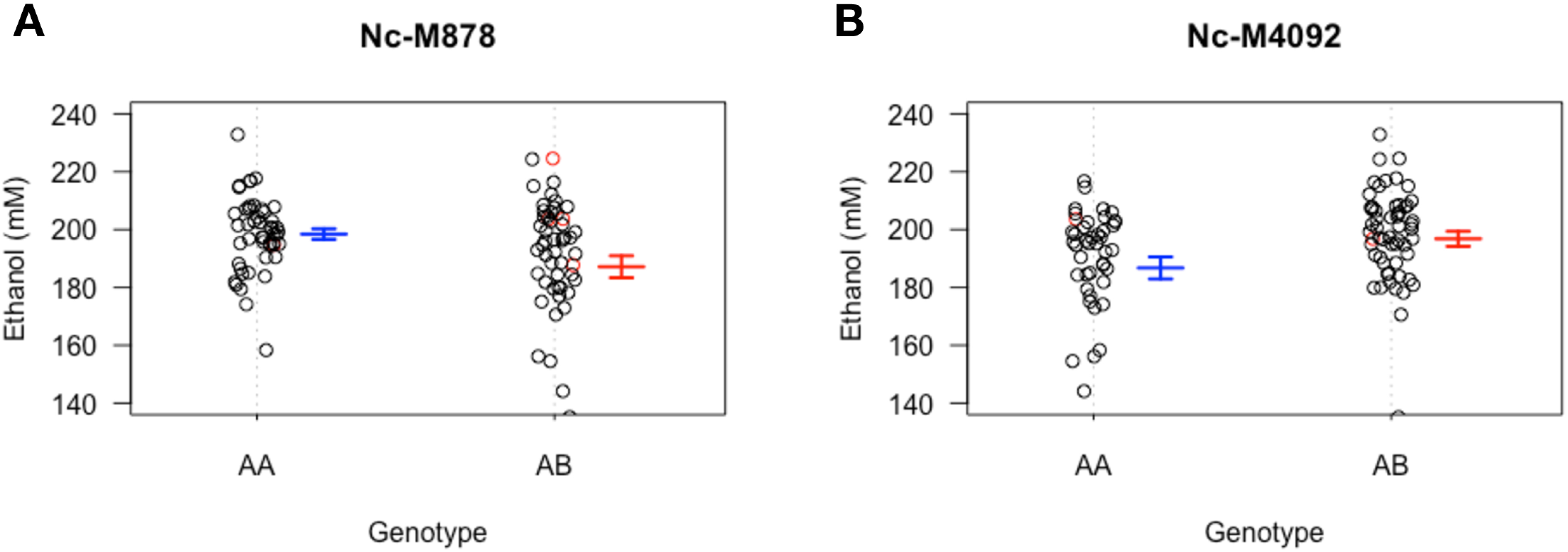
Phenotype Segregation at QTL Markers for Fermentation. A) Phenotype segregation at the QTL marker on LG I according to genotype. B) Phenotype segregation at the QTL marker on LG VI according to genotype. The error bars represent the error of the mean, n = 4 biological replicates. The data points in red are missing data at the given marker, however based on imputation it is inferred that they would be that genotype at the given marker and they are therefore plotted with that group.

Of particular interest in the linkage block in LG I, indicated by QTL analysis for fermentation, was a cluster of annotated glycolytic enzymes, sugar transporters, enzymes involved in alternate fates of pyruvate, including lactate dehydrogenase enzymes, and components of energy production machinery (Table 1). Alternatively, the only annotated gene of interest from the QTL on LG VI was another sugar transporter, *sut-7*.

**Table 1.** Candidate QTL genes for ethanol fermentation.

| Gene ID | Gene Name | Description | Linkage Group | Location |
| --- | --- | --- | --- | --- |
| NCU00629 | emp-3 | 6-phosphofructokinase | I | CM002236: 7,764,287 - 7,768,767 (-) |
| NCU00575 | gki-2 | glucokinase-2 | I | CM002236: 7,953,683 - 7,958,506 (-) |
| NCU00742 | glp-1 | glycerol-3-phosphate dehydrogenase | I | CM002236: 7,360,766 - 7,362,918 (+) |
| NCU00720 | tca-17 | L-lactate dehydrogenase | I | CM002236: 7,427,034 - 7,428,944 (+) |
| NCU00775 | tca-6 | isocitrate dehydrogenase subunit 1 | I | CM002236: 7,214,344 - 7,217,183 (-) |
| NCU00780 | ldh-4 | lactate dehydrogenase-4 | I | CM002236: 7,202,837 - 7,204,284 (+) |
| NCU00606 | null | short-chain alcohol dehydrogenase | I | CM002236: 7,858,582 - 7,859,522 (-) |
| NCU00801 | cdt-1 | MFS lactose permease, Sugar (and other) transporter | I | CM002236: 7,121,829 - 7,125,021 (-) |
| NCU00821 | sut-15 | sugar transporter-15 | I | CM002236: 7,056,000 - 7,058,684 (-) |
| NCU00809 | sut-26 | sugar transporter-26 | I | CM002236: 7,088,949 - 7,091,552 (+) |
| NCU05627 | sut-7 | sugar transporter-7 | VI | CM002241: 1,275,879 - 1,278,130 (-) |

Similarly, multiple QTL were observed for saccharification on LG II (Nc-M1321 = CM002240: 1924972: 184 Genes - 84 Annotated, 64 Hypothetical, 36 Unspecified), LG IV (Nc-M2984 = CM002239: 4860803: 184 Genes - 74 Annotated, 72 Hypothetical, 38 Unspecified), and LG VI (Nc-M4097 = CM002241: 1342332: 144 Genes – 44 Annotated, 54 Hypothetical, 16 Unspecified), although none were above the LOD threshold for 90% CI (Fig 5). Marker Nc-M1321 on LG II lies within an intergenic region, while markers Nc-M2984 on LG IV lies within a hypothetical protein (NCU07004) and marker Nc-M4097 on LG VI lies within phosphatidylinositol 3-kinase tor2 (NCU05608). Combined, the putative QTL account for 19.5% of the total variation observed in saccharification of cellulose. The major QTL on LG II accounts for 11% of the observed variation, while minor QTL on LG IV and VI account for 4.6% and 5.7%, respectively.

**Figure 5.**
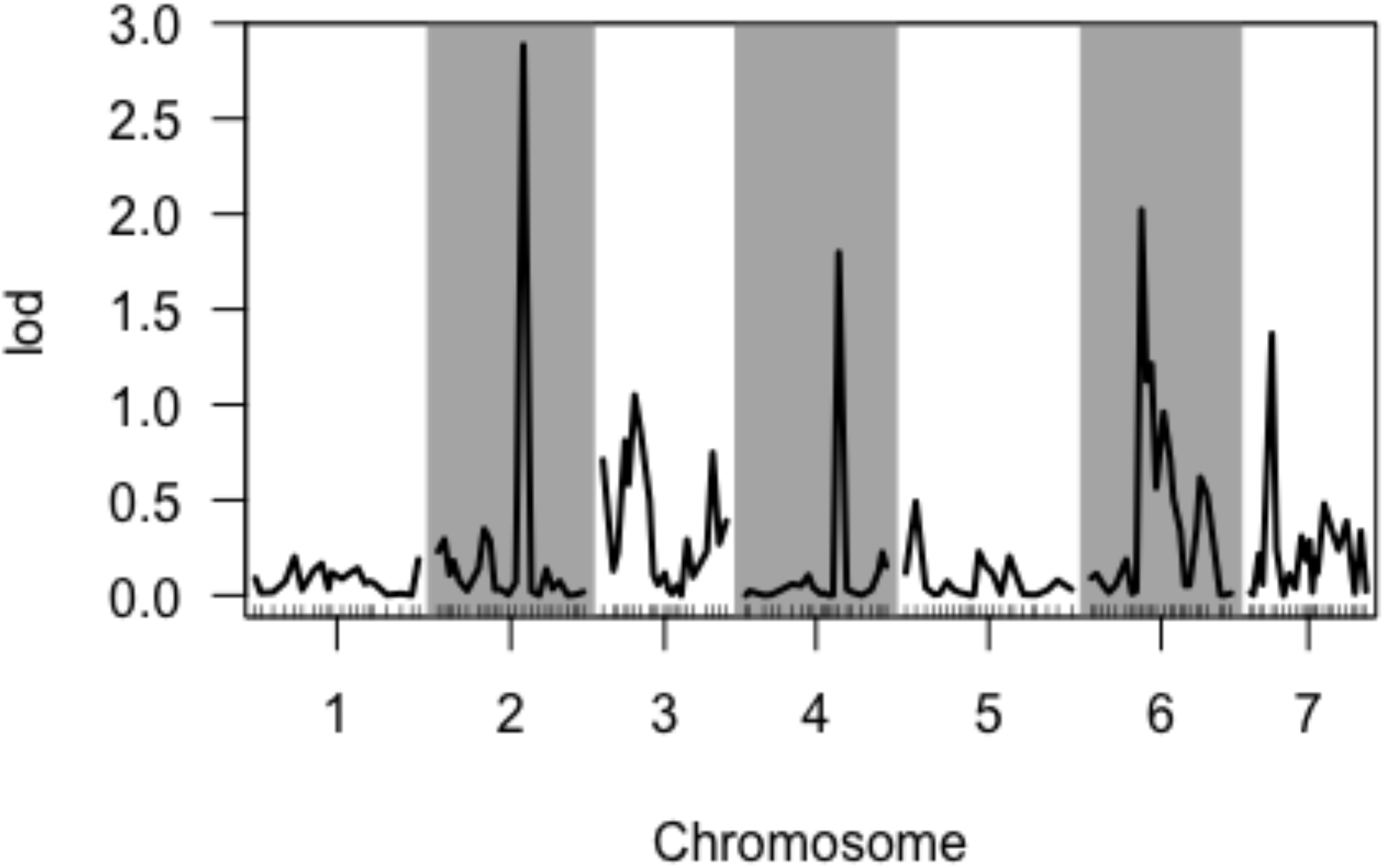
QTL analysis of Saccharification of Cellulose by the N6 population. Composite Interval Mapping single-QTL scan for FPA assay using linear regression (EM Algorithm). The y-axis is a measure of the logarithm of the odds ratio (LOD). Tick marks along the x-axis delineate relative marker positions, while alternating grey bands denote different chromosomes.

Marker segregation analysis was performed to confirm the estimation of variance analysis; which showed that the QTL in Chr2 contributed 11% of the variance (ANOVA F=14.647, p = 0.0002), while the one in Chr4 only contributed 4.6% variance (ANOVA F=6.078, p = 0.015) (Fig 6A and B). From this finding, we chose to investigate the potential for epistatic interactions among the QTL markers for saccharification of cellulose using R/QTL’s Scantwo function. Scantwo analysis revealed that the QTL marker on LG II (Nc-M1321) exhibited strong interactions with markers across all chromosomes, with the strongest interactions with the adjacent marker on LG II and the putative QTL marker on LG IV (Nc-M2984) (Fig 7A). Marker interaction plots demonstrated that the genotype B specific increases in FPA activity only held for the interactions between QTL markers on LG II and LG IV, but not with the interactions between adjacent markers on LG II (Fig 7B and C).

**Figure 6.**
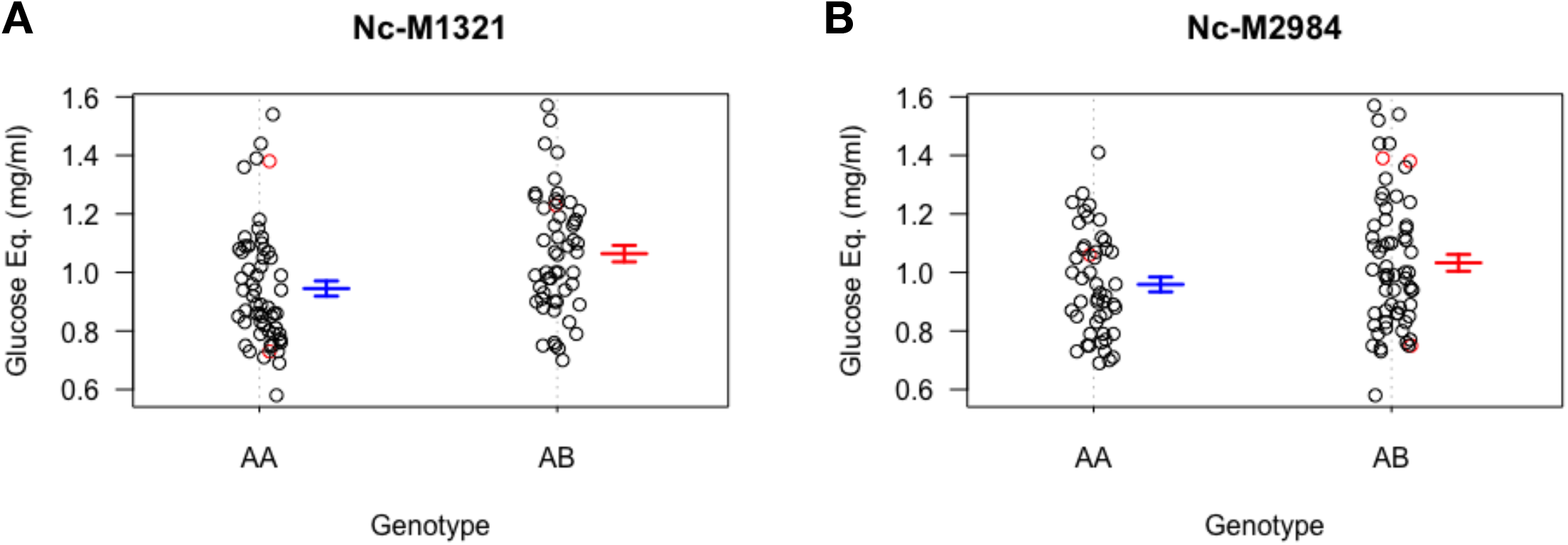
Phenotype Segregation at QTL Markers for Saccharification. A) Phenotype segregation at the QTL marker on LG II according to genotype. B) Phenotype segregation at the QTL marker on LG IV according to genotype. The error bars represent the error of the mean, n = 3 biological replicates. The data points in red are missing data at the given marker, however based on imputation it is inferred that they would be that genotype at the given marker and they are therefore plotted with that group.

**Figure 7.**
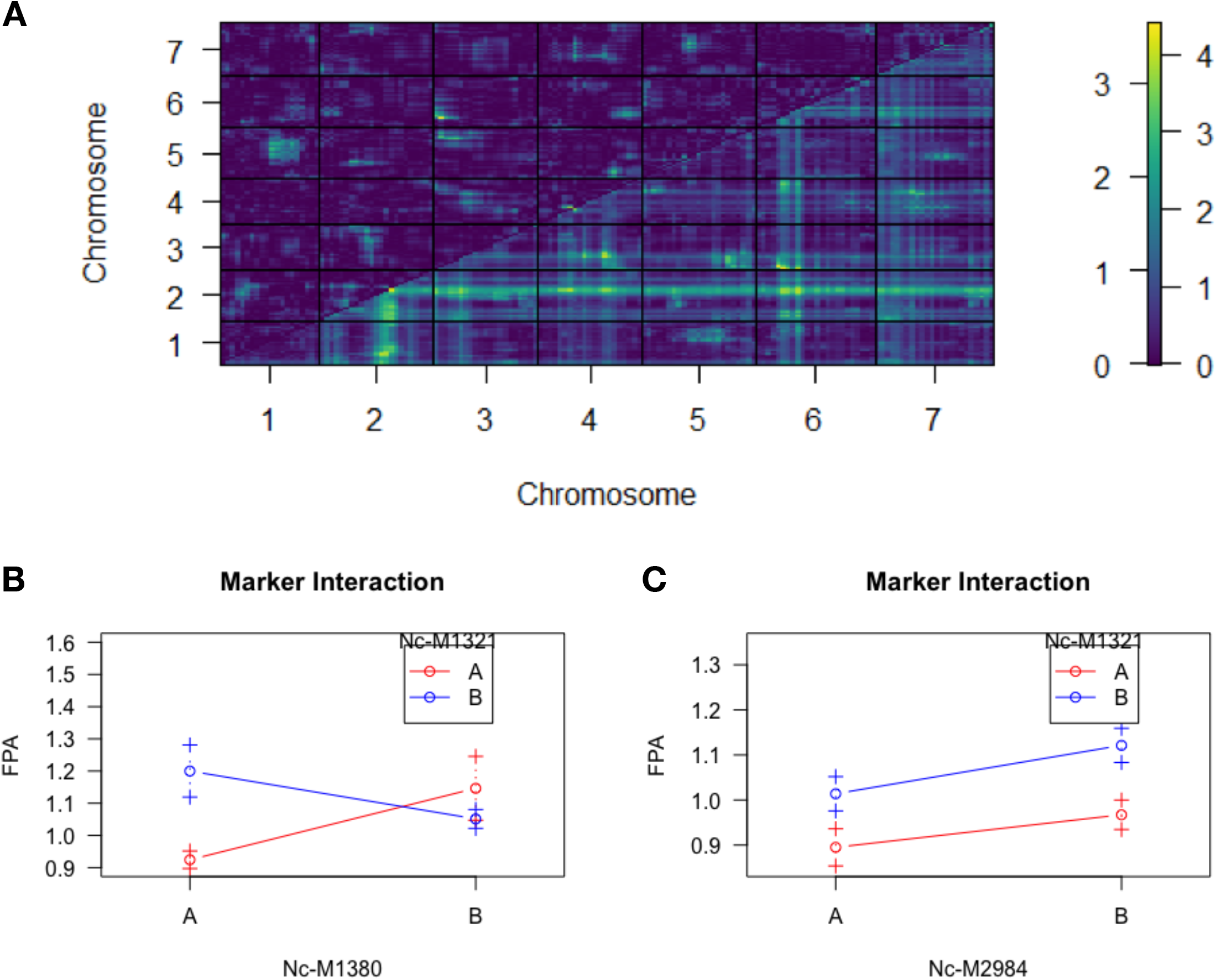
Saccharification QTL Marker Interactions. (A) Scantwo QTL analysis for epistatic marker interactions. Heatmap legend represents logarithm of the odds ratio (LOD) scores for marker interactions. (B) Average phenotypes according to genotype at sequential markers on LG II [Marker Nc-1321 is QTL marker on LG II]. (C) Average phenotypes according to genotype at QTL markers on LG II and LG IV. Colors represent genotype at marker indicated in legend, while letters on x-axis indicate genotype at marker indicated on x-axis title. Error bars represent standard deviations.

Among those genes within the QTL regions for saccharification there were some glucanase and xylanase enzymes, and transcription factors, which may have a role in sensing or degrading cellulose. However, two major regulatory genes of interest were also identified, *vib-1* and *xlr-1*, on LG II and LG IV respectively (Table 2).

**Table 2.** Candidate QTL genes involved in saccharification of cellulose.

| Gene ID | Gene Name | Description | Linkage Group | Location |
| --- | --- | --- | --- | --- |
| NCU03725 | vib-1 | VIB-1, variant 1 | II | CM002240: 2,215,819 - 2,219,221 (+) |
| NCU06971 | xlr-1 | transcriptional activator xlnR | IV | CM002239: 4,706,112 - 4,709,560 (-) |
| NCU01050 | gh61-4 | endoglucanase II | II | CM002240: 2,769,232 - 2,770,004 (+) |
| NCU03641 | gh3-1 | Beta-glucosidase 2, variant 1 | II | CM002240: 1,920,114 - 1,923,804 (+) |
| NCU01059 | gh47-3 | glycosyl hydrolase family 47 protein | II | CM002240: 2,790,429 - 2,792,645 (-) |
| NCU01050 | gh61-4 | endoglucanase II | II | CM002240: 2,769,232 - 2,770,004 (+) |
| NCU03657 | gh63-1 | mannosyl-oligosaccharide glucosidase, variant 1 | II | CM002240: 1,992,478 - 1,995,662 (+) |
| NCU01080 | gh64-1 | glucanase B | II | CM002240: 2,848,665 - 2,850,629 (-) |
| NCU06961 | gh28-2 | Exopolygalacturonase | IV | CM002239: 4,669,933 - 4,671,737 (+) |
| NCU07326 | gh43-6 | hypothetical protein | IV | CM002239: 4,359,254 - 4,361,686 (-) |
| NCU07005 | gh76-9 | glycosyl hydrolase family 76-9 | IV | CM002239: 4,865,028 - 4,867,282 (-) |
| NCU07269 | gh92-2 | alpha-1,2-mannosidase | IV | CM002239: 4,569,439 - 4,572,146 (-) |

## Discussion

In current study, we have identified that there exists a substantial variation in bioethanol-related traits in the mapping population, N6. We also identified a few major candidate genes with the potential to contribute to the observed variation, including the clustered glucose metabolism genes on LG I for fermentation, and transcriptional regulators *vib-1* and *xlr-1* on LG II and LG IV, respectively, for saccharification. The clustering of glucose metabolism genes on LG I makes identification of candidate genes for further analysis more complicated, requiring gene expression analysis for identification of those genes within the region that are differentially regulated and contributing to the observed variation in ethanol fermentation potential. While it is possible that an increased rate of glycolysis is responsible for increased fermentative capacity, the presence of lactate dehydrogenase enzymes within the QTL regions could present an alternative answer, with decreased lactic acid fermentation resulting in increased rates of ethanol fermentation.

The presence of major metabolic regulators within QTL regions for saccharification provide more promising QTL candidate genes. The first major gene of interest for saccharification of cellulose is the transcription factor, vegetative incompatibility blocked-1 (*vib-1*), located on LG II, which is involved in sensing carbon starvation and signaling for induction of the cellulolytic response (28). The second major gene of interest for saccharification of cellulose indicated is the xylose degradation regulator (*xlr-1*) located on LG IV, as it is involved in induction of the hemicellulose response, and regulation of cellodextrin transporters and cellulose degradation regulator *clr-1*, along with *vib-1*, identified in the putative QTL on LG II (12, 29).

Although *xlr-1* has not been shown to directly bind to promoters for cellulase enzymes, it has been implicated in cellulase production. Transcriptional analyses demonstrated that *xlr-1* knockouts have reduced hemicellulase and cellulase production when grown in xylan inducing conditions (12, 17). ChIP-seq experiments further revealed that although *xlr-1* does not directly bind to cellulase promoters it does bind to the promoter of transcription factor *clr-1*, a major regulator of cellulase expression, and can be found bound to targets in conjunction with *clr-1* and/or *clr-2* (12). Furthermore, *xlr-1* is involved in the regulation of *vib-1*, which in turn further regulates expression of *clr-1* (12). Similarly, Cai et al., demonstrated that *xlr-1*, not *clr-1/2*, is required for expression of cellodextrin transporter-2 (*cdt-2*), which is involved in sensing cellulose in the environment as well as transmembrane transport (30). Although cdt-2 was mildly reduced in the clr-1 knockout, cdt-2 expression was completely abolished with the deletion of xlr-1, suggesting that that xlr-1 is the main positive regulator of cdt-2 in N. crassa (27). *Cdt-2* has been implicated as a major cellodextrin importer for facilitating induction of the cellulase response, as cellulase production and growth on cellulose are significantly reduced in *cdt-2* knockouts (30, 31). Overexpression of *cdt-2* has also been shown to increase cellulase and hemicellulase production under cellulose and xylan conditions (30). These data could suggest pleiotropic or epistatic roles for *xlr-1* as a key component for cellulase production through its role in regulating of *vib-1* and *cdt-2* for carbon sensing and cellulase induction, *clr-1* for inducing the cellulolytic response via *clr-2*, and the co-regulation of genes targeted by *clr-1/2*. However, further experiments would be needed to address this.

Among other filamentous fungi, *xlr-1* orthologues (*xlnR/xyr-1*) have been shown to directly regulate cellulase expression alongside hemicellulase, although this role has been lost in *Neurospora* (12, 15, 17, 18, 32). However, a vestige of this role may be evident in *xlr-1*’s indirect role in the response to cellulose through its regulation of critical genes involved in sensing cellulose in the environment, especially through *vib-1, clr-1, cdt-2* and major facilitator superfamily (MFS) sugar transporters. While RNA-seq and ChIP-seq experiments have previously been employed to decipher the role of *xlr-1* in *N. crassa*, these studies have used knockout and overexpression strains that share the same genetic scaffold as the wild type reference. Similar studies using diverse natural strains may demonstrate that differential expression of *xlr-1* leads to differences in cellulolytic potential or illustrate potentially disparate roles arising from allelic variation at the *xlr-1* locus or its targets.

Allelic variation in lignocellulose degradation regulators and their target genes may provide unique insights into the lignocellulose response of *N. crassa*. Transcriptomic profiling and expression analysis may reveal differential interactions between *xlr-1* and its’ targets, resulting in differential enzyme induction and degradation of cellulosic substrates. Furthermore, expression QTL (eQTL) analysis would be important for discerning the pleiotropic effects of *xlr-1* from its epistatic effects. Understanding the implications of allelic effects on lignocellulose metabolism should allude to elite genotypes for further enhancement and possible industrial applications in bioethanol production.

## Acknowledgements

The authors thank Drs. Sunil Shende and Charot Rodeget for curating Python scripts, critical reading and insightful comments.

## Author Contributions

**Conceptualization:** Kwangwon Lee and Joshua C. Waters

**Project Administration:** Kwangwon Lee

**Funding Acquisition:** Kwangwon Lee

**Methodology:** Joshua C. Waters and Kwangwon Lee

**Supervision:** Joshua C. Waters and Kwangwon Lee

**Investigation:** Joshua C. Waters, Deval Jhaveri, Justin C. Biffinger

**Validation:** Joshua C. Waters, Deval Jhaveri, Justin C. Biffinger, Kwangwon Lee

**Data curation:** Deval Jhaveri

**Software:** Joshua C. Waters

**Formal Analysis:** Joshua C. Waters

**Visualization:** Joshua C. Waters

**Writing-original draft:** Joshua C. Waters and Kwangwon Lee

**Writing-review and editing:** Joshua C. Waters and Kwangwon Lee

## Supporting Information

**S1 File. N6_QTL_Data.** This .csv file contains genotype and phenotype information used for QTL analysis using the R-qtl package in R.

**S2 Text. N6 QTL script for analysis in R.** This file contains R script used for QTL analysis in R. The numbers in row two is a chromosome number and row 3 a physical base-pair position.

**S3 File. GenesByLocation_summary (Chr1-Ferm).xlsx.** This file contains all of the genes within the QTL region on chromosome 1 for fermentation.

**S4_GenesByLocation_summary (Chr2-Sacch).xlsx.** This file contains all of the genes within the QTL region on chromosome 2 for saccharification.

**S5_GenesByLocation_summary (Chr4-Ferm).xlsx.** This file contains all of the genes within the QTL region on chromosome 4 for fermentation.

**S6_GenesByLocation_summary (Chr4-Sacch).xlsx.** This file contains all of the genes within the QTL region on chromosome 4 for saccharification.

**S7_GenesByLocation_summary (Chr6-Ferm).xlsx.** This file contains all of the genes within the QTL region on chromosome 6 for fermentation.

**S8_GenesByLocation_summary (Chr6-Sacch).xlsx.** This file contains all of the genes within the QTL region on chromosome 6 for saccharification.

**S9_GenesByLocation_summary (Chr7-Ferm).xlsx.** This file contains all of the genes within the QTL region on chromosome 7 for fermentation.

